# Intellectual contributions meriting authorship: Survey results from the top cited authors across all science categories

**DOI:** 10.1101/323519

**Authors:** Gregory S. Patience, Federico Galli, Paul A. Patience, Daria C. Boffito

## Abstract

Authorship is the currency of an academic career for which the number of papers researchers publish demonstrates creativity, productivity, and impact. To discourage coercive authorship practices and inflated publication records, journals require authors to affirm and detail their intellectual contributions but this strategy has been unsuccessful as authorship lists continue to grow. Here, we surveyed close to 6000 of the top cited authors in all science categories with a list of 25 research activities that we adapted from the National Institutes of Health (NIH) authorship guidelines. Responses varied widely from individuals in the same discipline, same level of experience, and the same geographic region. Most researchers agreed with the NIH criteria and grant authorship to individuals that draft the manuscript, analyze and interpret data, and propose ideas. However, thousands of the researchers also value supervision and contributing comments to the manuscript whereas the NIH recommends discounting these activities when attributing authorship. People value the minutiae of research beyond writing and data reduction: researchers in the humanities value it less than those in pure and applied sciences; individuals from Far East Asia and Middle East and Northern Africa value these activities more than anglophones and northern Europeans. While developing national and international collaborations, researchers must recognize differences in peoples values while assigning authorship.

## Introduction

The scientific process requires ingenuity and individuals that contribute creativity to answering the research question merit authorship [1, 2]. However authorship lists continue to climb [3, 4] despite the widespread dissemination of guidelines to dissuade ambiguous attribution [5, 6]: 5 to 10 individuals co-author work in 2013 that had only 2 to 3 in 1993 [7, 4]. The International Committee of Medical Journal Editors (ICMJE) and other organizations [8, 9] published a list of criteria for authorship requiring authors to: (1) design experiments, or analyse data, or interpret data; (2) draft or revise the manuscript; (3) approve the final manuscript; and, (4) agree to be held accountable for it [10, 11].

> In addition to being accountable for the parts of the work he or she has done, an author should be able to identify which co-authors are responsible for specific other parts of the work.

Furthermore, they emphasize that any individual that contributed to the first activity have the opportunity to participate in activities 2, 3, or 4, so as not to exclude them from authorship. The WAME criteria [8] are similar but somewhat broader and explicitly state that it is dishonest to disregard individuals that contribute to writing (ghost authorship) particularly those working for commercial companies. Like ICMJE they identify activities that warrant an acknowledgement:

> Performing technical services, translating text, identifying patients for study, supplying materials, and providing funding or administrative oversight over facilities where the work was done are not, in themselves, sufficient for authorship, although these contributions may be acknowledged in the manuscript, as described below. It is dishonest to include authors only because of their reputation, position of authority, or friendship (guest authorship).

Whereas the IJCME require all authors to be familiar with all aspects of the work and capable of identifying who did what, the WAME criteria recognize that a biostatistician contributes complementary expertise and may be incapable of defending all the clinical aspects of the work. Harvard Medical School adapt these criteria and insist that specialized personnel be included even when their contribution is limited in scope. One individual of the team takes responsibility for the document and keeps a record of how everyone contributed [12], like the WAME guarantor or the corresponding author (CA) for the ICMJE.

The National Institutes of Health (NIH) expanded the ICMJE criteria and defined 15 activities related to publishing research [4, 23]. They discourage honorary authorship and recommend: excluding individuals that train/educate, provide resources, or read/comment the manuscript; always including researchers that draft the manuscript or perform original experimental work; and include individuals that participate in the other activities depending on their implication. Earlier critiques of the ICMJE guidelines alleged that they created orphan papers in which nobody met all 4 criteria and thus no one was eligible to be considered as an author [24] but this oversight has been corrected.

Authorship criteria ain to reduce unethical practices—coercive authors, honorary authorship, guest authorship, gift authorship and ghost authorship. A cross-sectional survey of corresponding authors publishing in 6 high impact biomedical journals confirmed the guidelines reduced ambiguous authorship from 29 % in 1996 to 21 % in 2008 but it was mostly due to reducing ghost authorship [5]. The survey identified 17 functions related to developing an article and asked how many co-authors contributed to only one, which would make them ineligible for authorship according to ICMJE [13]. Compliance to these criteria in ecological research is much lower [14] where 78 % of the studies had at least one co-author that failed to meet ICMJE guidelines. Another study showed that in the top 1 % of the highest cited articles across 22 Web of Science Core Collection (WoS) journal fields, one-third of them included a specialized author and one-half with non-author collaborators (ghost authors) [15].

Researchers disregard the guidelines because they are too restrictive as they discount the minutiae of research and the *plurality of value and the messiness of scientific practice* [16]. Moffat (2018) [17] argues that a universal consensus of assigning authorship is neither attainable nor desirable because it infringes on the autonomy of researchers scientific expression. Knowing whose suggestion, insight, or initiative contributed (and by how much) to research success is unknowable *a priori* and even *a posteriori*. Furthermore, comprehensive criteria must recognize that theoreticians value knowledge and writing as core activities while applied scientists and engineers value data reduction, maintaining, designing, and operating equipment and other tools more. Consider *Big Science* that tackles challenges facing society with hundreds and thousands of researchers working as a group. Experimental high energy physics articles approach 3000 individuals routinely [18, 19] and the record for the most authors is 5154 [20]. One quarter of the top 500 cited articles in nuclear physics averaged 1160 authors (WoS, 2010 to 2015) [21]. Author counts biomedical journals are not so high but 19 of the 244 articles *Lancet* publsihed in2017 had more than 40 authors, 10 had more than 480 authors, and one had 1039 [22].

## Materials and methods

We expanded the NIH authorship activity list to include 25 research tasks and developed a questionnaire to gauge the practices of researchers across all scientific endeavors. We first developed the survey in Excel for a conference and refined the questions after feedback from students and colleagues [26]. In the following three months, we sent emails with a link to a our refined questionnaire with the MonkeySurvey platform to students and staff at Polytechnique Montréal, colleagues from other institutes, and companies. We then posted the link on Facebook and LinkedIn. Approximately 400 people responded, most of whom worked in chemical sciences in Iran, Canada, and Italy. Half of these respondents were senior professionals/professors and the rest were graduate students, researchers with less than 5 years experience, and business people. Our next mailing list included researchers in various scientific fields from 15 institutes in the United States, Great Britain, France, Germany, Singapore, and Japan. Approximately 60 individuals completed the survey: 30 in the first mailing and another 30 after a reminder.

We revised the questionnaire a final time stating that N/A (not applicable) was the same as not responding. In October 2017 we sent emails to corresponding authors of the top 500 cited articles from 2010 to 2014 in 235 WoS categories [21]. The email mentioned the title of the CA’s paper, its rank within the scientific category, and the total number of papers in the category. It stated that the survey took 5 minutes to complete, had 5 categories with 5 questions each. Approximately 84 000 researchers received the email and 3500 responded while 30 000 were returend undelivered.

A follow-up email included a link to bibliometric data of the 500 top cited articles in the CA’s scientific category. A further 3000 researchers attempted the survey at a completion rate of 91 %. The overall response rate was almost 10 %, which is less than half of the respondents in an earlier study related to contribution statements [25].

The five groups of questions resemble the classes of activities in journal contribution statements—supervision (conception), design (materials), execution, data reduction (analysis), and writing [25] with five activities per category. For each activity, respondents were to choose one of five options that corresponded to how often they thought it merited authorship (rather than how they thought others did it). We assigned a score *ω_i_* from 0 to 4 for each:

*ω*_0_ = 0: almost never—< 5 % of the time;
*ω*_1_ = 1: sometimes—25 % of the time;
*ω*_2_ = 2: often—50 % of the time;
*ω*_3_ = 3: usually—75 % of the time; and,
*ω*_4_ = 4: almost always—> 95 % of the time.

This scoring scheme resembles a statistical distribution where *ω*_2_ = 2 represents the mean; *ω*_4_=4 represents the 95 % confidence level and is 2 *σ* greater than the mean; *ω*_0_ = 0 is at the 5 % confidence level so it is 2 *σ* less than the mean; and finally, *ω*_3_ loosely represents 1 *σ* greater than the mean and *ω*_1_ represents 1 *σ* less than the mean.

The mean score of activity *i*, *s̄*_*i*_, corresponds to the quotient of the sum of the responses and the total number of responses:*s̄*_*i*_ = Σ*ω*_*i*,*k*_/*n*_*i*_. When *s̄*_*i*_ > 3, the overwhelming consensus confirms that activity *i* merits authorship. When *s̄*_*i*_ < 1, most people think that the activity *i* seldom if ever, merits authorship. However, even for the activities where *s̄*_*i*_ < 1, over 1000 individuals chose usually or almost always. The aggregate individual score (indice *j*) *S*_*j*_ = Σ_q=1,25_ω_*jq*_ varies from 0 to 100 and we use this metric to compare responses across fields, geography, experience, and work place.

### SurveyMonkey questionnaire

#### Supervision

Supervision includes preparing grants, mentoring subordinates, and securing funding but does the time dedicated to these activities constitute sufficient intellectual involvement to be considered an author? Together with approving the final manuscript, any one of the five activities merits authorship: almost never (< 5% of the time), sometimes (25 %), often (50 %), usually (75 %), or almost always (>95%).

**Q1** Securing funding
**Q2** Establishing the team
**Q3** Coordinating tests
**Q4** Proposing ideas
**Q5** Providing resources (laboratory space, analytical, time)

#### Experimental design (equipment)

Designing, operating, and maintaining experimental equipment are essential to generate data. Together with approving the final manuscript, any one of the five activities merits authorship: almost never (< 5% of the time), sometimes (25 %), often (50 %), usually (75 %), or almost always (>95%).

**Q6** Setting up experimental equipment, writing programs
**Q7** Designing equipment, writing programs
**Q8** Operating instruments and equipment, running programs
**Q9** Modifying, maintaining equipment and programs
**Q10** Troubleshooting mechanical failures

#### Sample manipulation

Researchers generate, analyze and share samples with others for further analysis. Together with approving the final manuscript, any one of the five activities merits authorship: almost never (< 5% of the time), sometimes (25 %), often (50 %), usually (75 %), or almost always (> 95%).

**Q11** Identifying necessary samples for the program
**Q12** Generating samples for external analysis (by collaborators or third parties)
**Q13** Supplying samples, computer programs
**Q14** Analysis of samples by third parties (that you pay for)
**Q15** Discuss results of samples, viability, reliability, error

#### Data reduction

Together with approving the final manuscript, any one of the five activities related to manipulating/analyzing data merits authorship: almost never (< 5% of the time), sometimes (25 %), often (50 %), usually (75 %), or almost always (> 95%).

**Q16** Developing an experimental plan (DOE)
**Q17** Collecting/measuring experimental data, executing programs
**Q18** Consolidating experimental data
**Q19** Analyzing data, identifying trends
**Q20** Interpreting results, modelling, deriving correlations

#### Writing

Writing papers includes adding text to sections, revising the document and responding to referees. Together with approving the final manuscript, any one of the five activities merits authorship: almost never (< 5% of the time), sometimes (25 %), often (50 %), usually (75 %), or almost always (> 95%).

**Q21** Major role in drafting document
**Q22** Commenting on scientific content
**Q23** Correcting language, grammar, sentence structure
**Q24** Proofreading and suggesting substantial modifications
**Q25** Responding to reviewers/editors comments

The questions remained succinct so as not to unduly influence the respondents and to minimize the time to do the survey. The mean response time was 5 minutes with a median of 4 minutes (Fig 1). Some commented that the questions were vague and thus open to interpretation but the email included references to an earlier study [4] with the NIH classification [23] and our contact information. Many referred to the article, few called, but we responded to 1000 messages.

**Fig 1.**
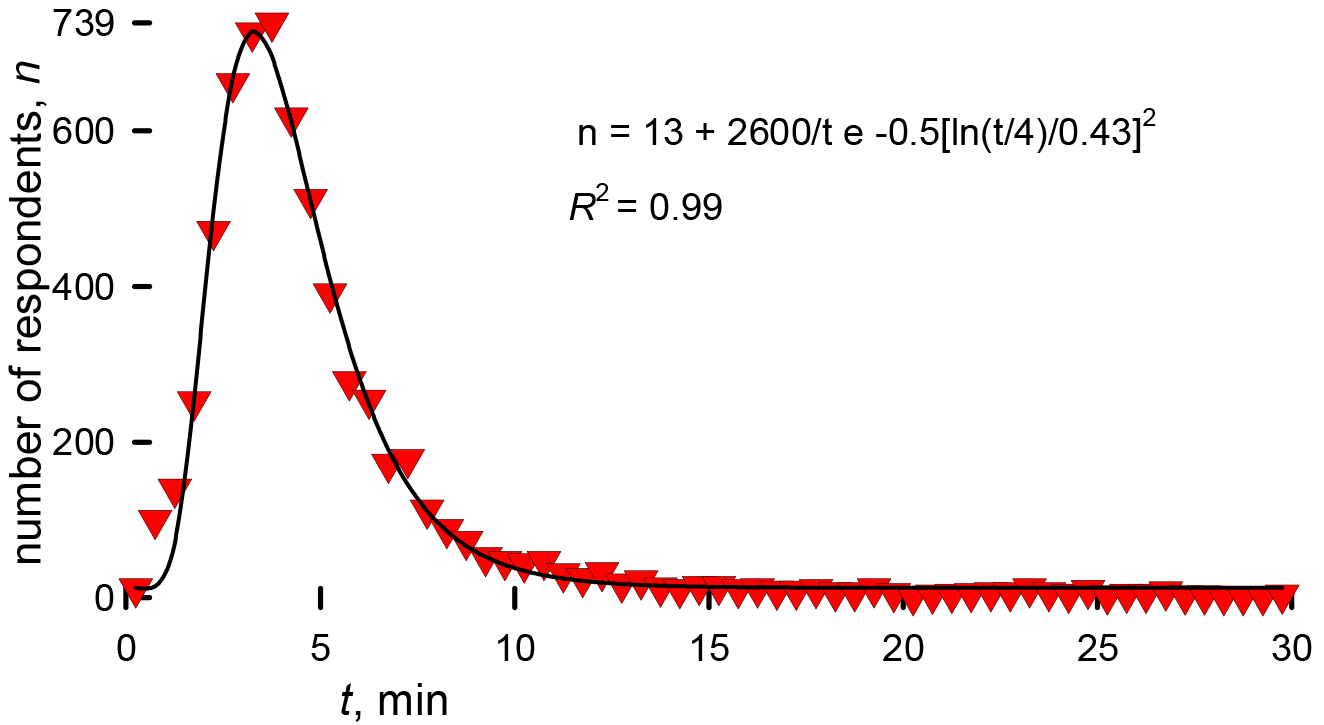
Time to complete the survey.

We retained 5781 responses of 6604 researchers that participated in the survey. We rejected responses that took less than one-minute to complete and those with a standard deviation equal to zero, which means all questions had the same response. We examined responses for which *S_j_* > 90 and *S_j_* < 10 and elminated those whose responses were incomplete (*n* < 10) or incoherent. The blue line in Fig 2 represents the the scores of all individuals and the bars corresponds to the retained responses. The standard deviation and mean for the reduced data set were 16 and 54.4, respectively, while for the entire data set they were 19 and 51.8.

**Fig 2.**
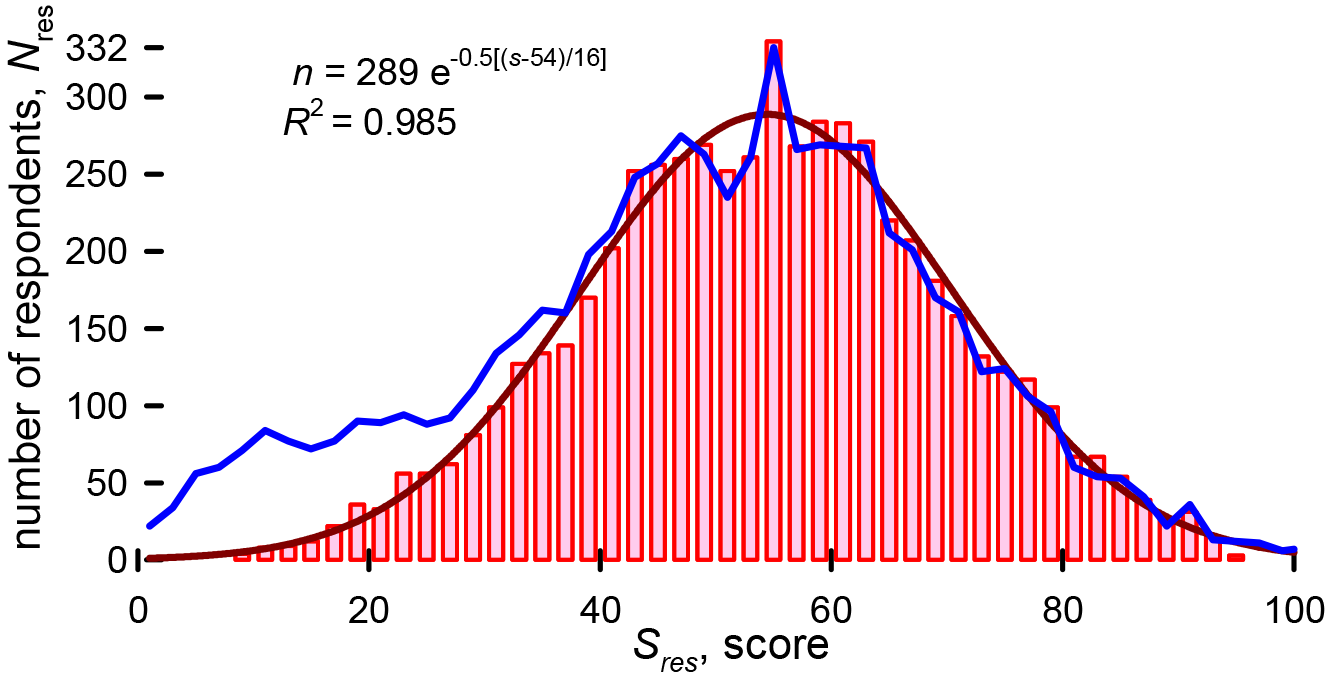
Frequency of individual scores: Blue line includes all responses, *N* = 6604, and an average score, *S*, of 48.6; bars represent retained responses, *N* = 5781 and an average score of 53.7; red line is the best first Gaussian distribution, *R*^2^ = 0.985.

Together with 25 questions related to research activities, we collected biographical data: seniority, country of birth, work place, and and research discipline.

**Q26** Identify your level of experience

Senior researcher/professor (> 5 y)
Early career professional/professor (< 5 y)
Senior graduate student (> 2 y)
Junior graduate student (< 2 y)

Most of the respondents had a least 5 years of professional experience (5129), followed by early career professionals with less than 5 years (467). Few students participated in the study (180 with more than 2 years and 83 with less than 2 years) and even fewer in management (53).

**Q27** What is your country of birth? Researchers born in 115 countries participated in the survey. Americans responded most (1296), followed by the British (452), Italians (369), Germans (359), and Canadians (339). The number of respondents from each country should correlate with the number of corresponding authors that wrote the most cited articles.

**Q28** Where do you work?

University
Government
Institution
Company
Other (please specify)

Most of the participants worked for universities (4214), followed by institutions (670), government (281) and finally companies (209). There were 206 individuals that chose *other* including: multiple affliations—academic and hospital (47), private-practice and consultants (30), hospitals, international, and national institutes (49), non-governemental agencies and museums (20).

**Table 1.**
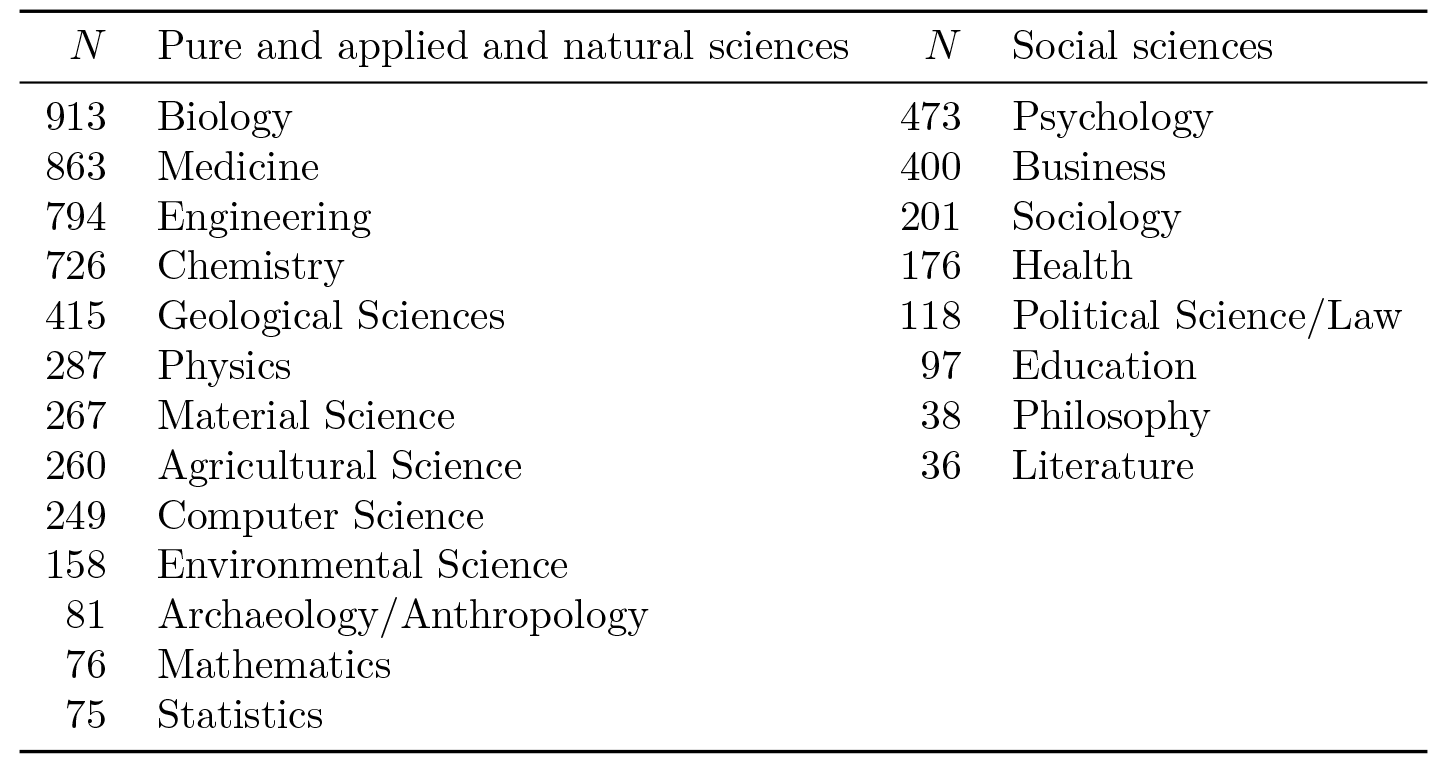
Participation rates based on scientific discipline allowing for multiple affiliations between categories (*N_T_* = 6703). The total number of independent responses was 5363.

**Q29** What is your research discipline? Over 1000 individuals cited multiple research disciplines like chemistry and physics, or behavioural neuroscience and psychology. Others referred to topics like molecular evolution, ultrasound, biomedical, and bone rather than specifying a category. We consolidated the answers and assigned each response to one of 22 scientific categories following closely the classification of Clarivate Analytics Science Watch [27] (Table 1). A total of 5363 reponded to this question but over 1000 included multiple responses so the total was 6703. The lowest numbers of respondents were in literature (36), philosophy (38), mathematics (76), and statistics (76). Researchers in the humanities, mathematics, theoretical physics, and economics stated that most of the questions did not apply to them. Individuals in biology, medicine, engineering, and chemistry responded with the highest frequency, which corresponds to the most categories in WoS (Table 1).

## Results

Most CAs agreed to assign authorship to those that drafted the manuscript (*s̄*_21_ = 3.7), interpreted data (*s̄*_20_ = 3.6), and analyzed data (*s̄*_19_ = 3.3), which agrees with the ICMJE criteria (Fig 3). However, unlike the ICMJE, they attributed authorship to many other activities like proposing ideas (*s̄*_19_ = 3.3), consolidate data (*s̄*_15_ = 2.7), execute DOE (*s̄*_15_ = 2.8), experimental design) (*s̄*_15_ = 2.8, and responding to reviewers (*s̄*_25_ = 2.7). Execute (sample management) and design (including operation) were the least valued categories, but even so, thousands of people thought that activities in these categories always merited authorship.

**Fig 3.**
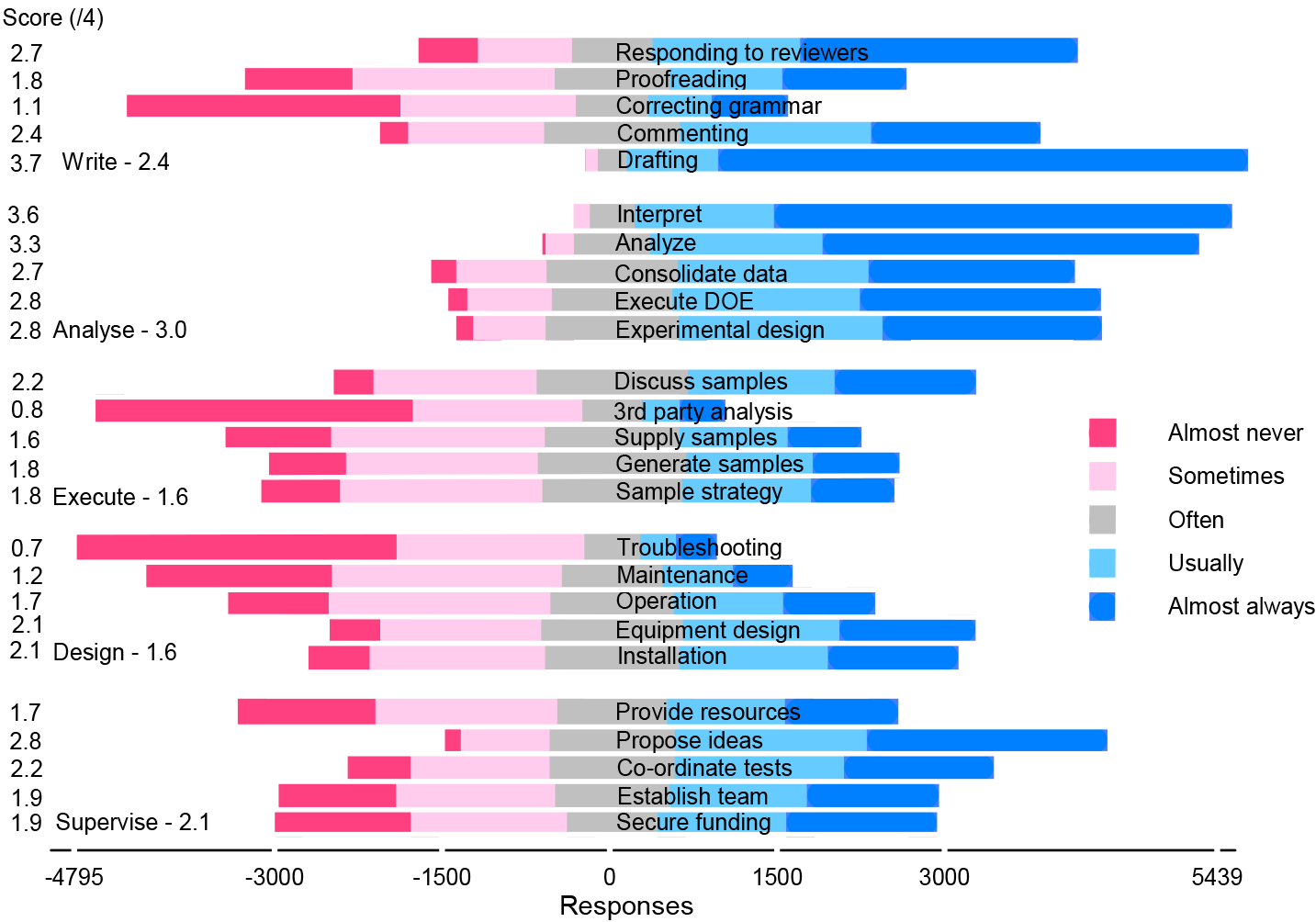
All fields Likert chart: 5500 respondents. The number of responses less than zero on the *x*-axis include almost never (*ω*_0_), sometimes (*ω*_1_) and half of the often (*ω*_2_) responses. The number of responses greater than zero include almost always (*ω*_4_), usually (*ω*_3_), and half of the often responses. The mean scores on the *y*-axis are *s̄_i_* = Σ_*k*=1,*n*_*i*__ ω_*i,k*_/*n*_*i*_ *s*̄_*i*_ and vary from a low of 0.7 (troubleshooting) to a high of 3.7 (drafting).

Researchers were most ambivalent about supervision: with a score of 2.1: more that one thousand thought these activities almost always merit authorship and about the same thought they almost never merit authorship. Proposing ideas scored 2.8 (*s̄*_4_), ranking it fourth among the 25 activities, while providing resources scored poorly at 1.7 (*s̄*_5_). Even so, one third of the respondents indicated that providing resources merits authorship almost always or usually.

The responses ranged from almost always to almost never for most activities independent of the scientific category, even for people in the same region, and same level of experience. However, to identify differenece in general tendencies, we calculated mean scores according to category (Fig 4). The standard deviation between category and means were highest for drafting the document and analyzing data. Scores diverged substantially for the other activities, although the trends were similar. The lowest repsonse rates were from philosophy and literature and they also had the lowest scores across most questions: they were most unlikely to grant authorship for anything other than writing and analysis. Researchers in materials sciences were more likely to grant authorship for other activities.

**Fig 4.**
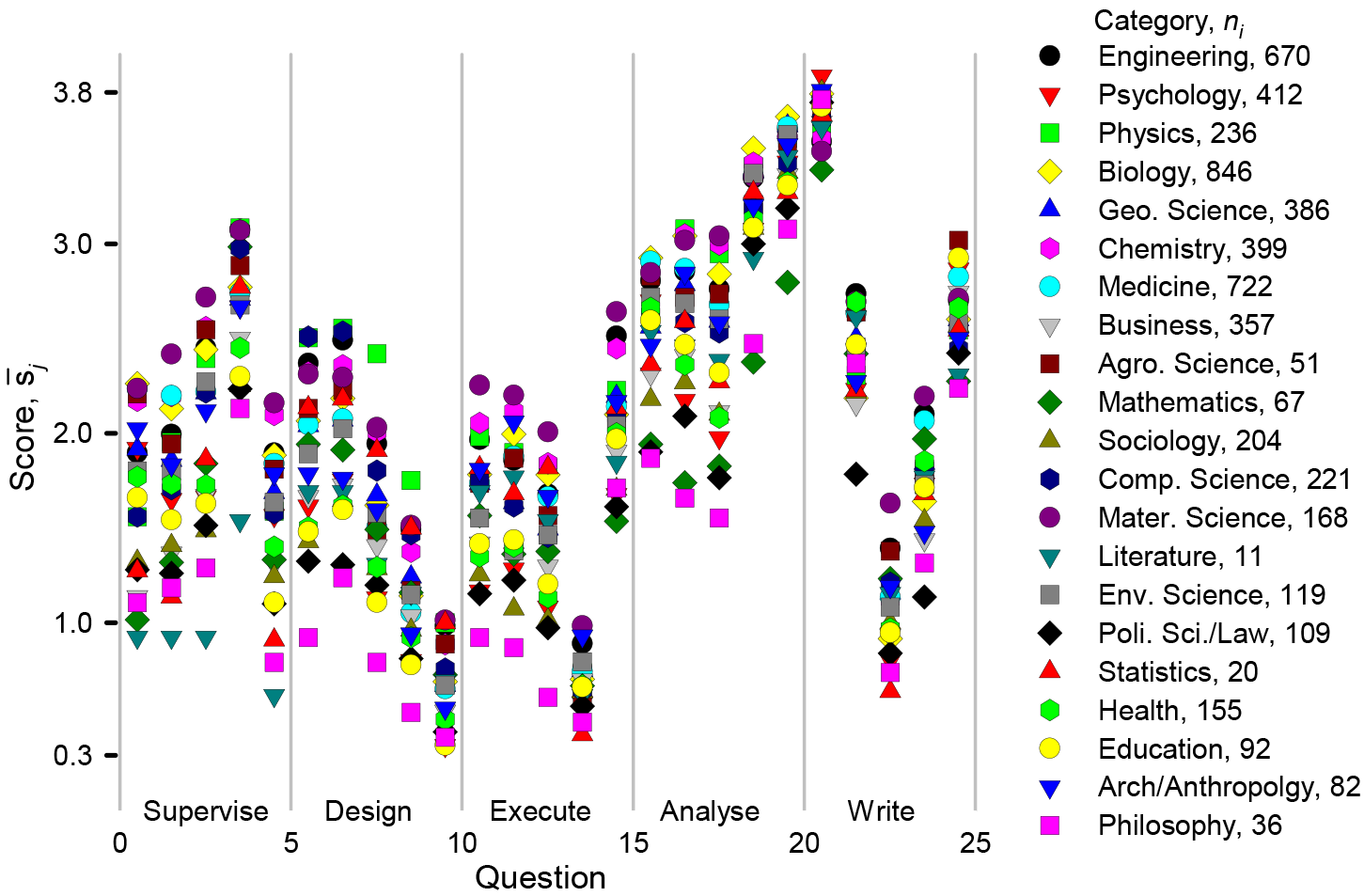
Mean scores,*s*̄_*j*_, according to scientific category. We grouped 5363 respondents according to the Clarivate Analytics science category definitions [27] and then calculated a mean score per category based on the total number of respondents in each category, *N*_*j*_: *s*̄_*i*_ = Σ_*k*_ω_*j,k*_/*N*_*i*_.

We grouped the categories into four broad fields—pure and applied sciences, natural science, humanities, and philosophy, law, and political science (Fig 5). The average score for the pure and applied sciences was *S̄* = 59 and it was lowest for philosophy, political science at *S̄* = 41. Differences between pure and applied sciences and natural sciences (*S̄* = 54) were about the same as that between natural sciences and humanities (*S̄* = 48). The size and scope of collaborative teams in the humanities is much lower than in pure and applied and natural sciences, which accounts for some of the difference in the scores [21].

**Fig 5.**
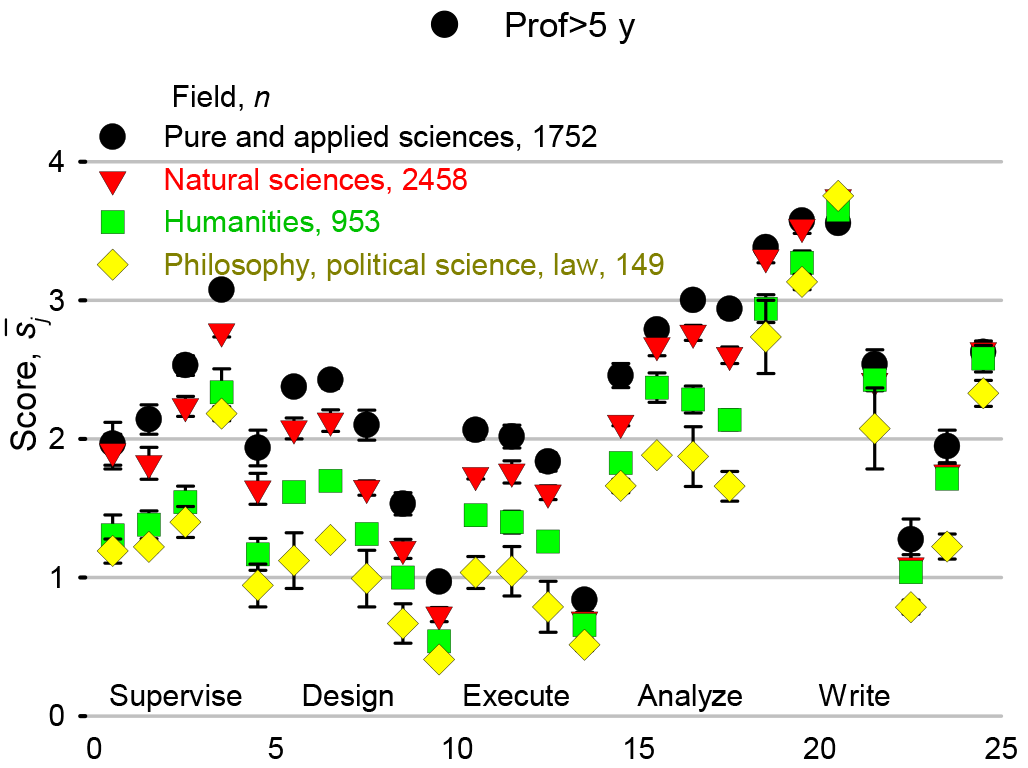
Mean scores according to scientific field. We compared the aggregate category scores, *S̄*, and assigned them to one of four fields based on the Student’s *t*-test. Pure and applied sciences includes physics, chemistry, materials, and engineering (*S̄* = 59). Natural sciences includes biology, geosciences, medicine, computer science, agricultural sciences, environment, statistics, and archaeology (*S̄* = 54). Many humanities categories made up the third field (*S̄* = 48): psychology, management, mathematics, sociology, literature, education, and health. The final field (*S̄* = 41) comprised philosophy, political science, and law.

To test whether opinions varied according to experience and birth country, we only considered pure and applied and natural sciences (column 1 of Table 1). We ordered aggregate scores from each question, *s*̄_*i*_ from the lowest to the highest(Fig 6). Responses of professionals with less than 5 years of experience and those with more than 5 years were highly correlated (*R*^2^ = 0.985). Graduate students (predominantly Master’s) had the second highest correlation with the professionals (*R*^2^ = 0.95). Graduate students with more than 2 years of experience viewed the contributions from each of the activities more favourably, while business people were less inclined to grant authorship to most of the activities. For all groups, the highest three and lowest three ranked activities were the same. Most thought that preparing a grant proposal would often merit authorship while business people thought that it would warrant authorship only sometimes.

**Fig 6.**
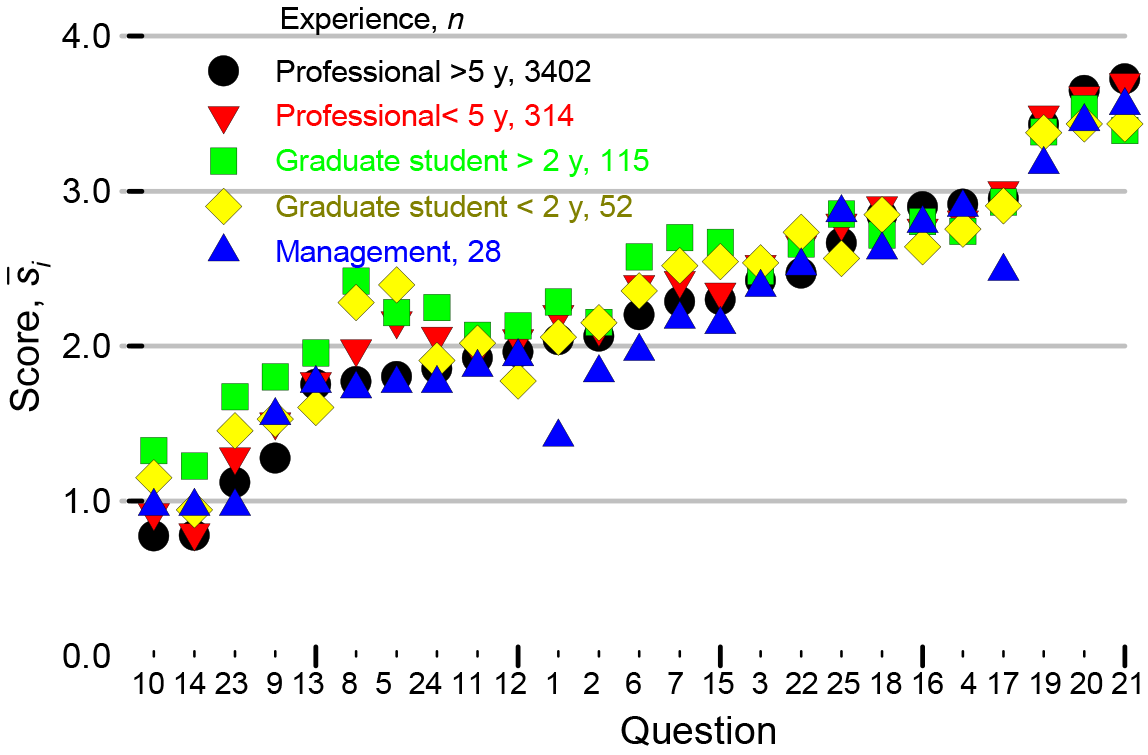
Scores for each of the questions with respect to career experience. The questions were organized from lowest to highest score based on the answers of the professionals with 5 years of experience.

To test tendencies based on linguistic and regional circumstances, we initially grouped the countries into 8 categories: (1) hispanophone, (2) anglophone (including Canada with a significant French population), (3) Eastern European (including ex-USSR states), (4) South Asia (including Iran, Afghanistan, and Pakistan), (5) Western Europe (and Israel), (6) sub-Saharan Africa, (7) Middle East and North Africa, (8) and Far East Asia. We then compared the means of responses from individual nations against the regions. Based on a *t*-test comparing the *s*̄_*i*_ of each country against the group average, we reformed the groups. Responses from Italians, French, Greek, and Cypriots resembled the hispanophones more than the rest of Europe, so we grouped them together and labelled them Latin (although Greek is not a Latin language). Northern Europe we labelled Germanic even though Finnish is not Germanic. We also labelled former Eastern European countries Slavic although Hungary, Romanian, and some of the Soviet Bloc states speak other languages. Because the sub-Saharan Africa group only had 22 individuals, we combined it with the Middle East and North Africa (MEA). We could have grouped these nations with anglophones as most are part of the Commonwealth. Based on the original *t*-test, Iran belonged with South Asia but with the expanded MEA grouping, it belongs to either South Asia or MEA, so we regrouped Iran with the latter.

The sample size for answers to countries was 5669 (Fig 7). Americans responded most and the anglophone group hsd twice as many respondents as the latin and germanic groups. Germany followed by the Netherlands headed the germanic group and the next 5 countries each had at least 50 respondents. The distribution for number of respondents per country for the latin group was similar with Italy heading it followed by France and Spain. The other regions had a similar representation that varied from 242(South Asia) to 385 (Far East Asia).

**Fig 7.**
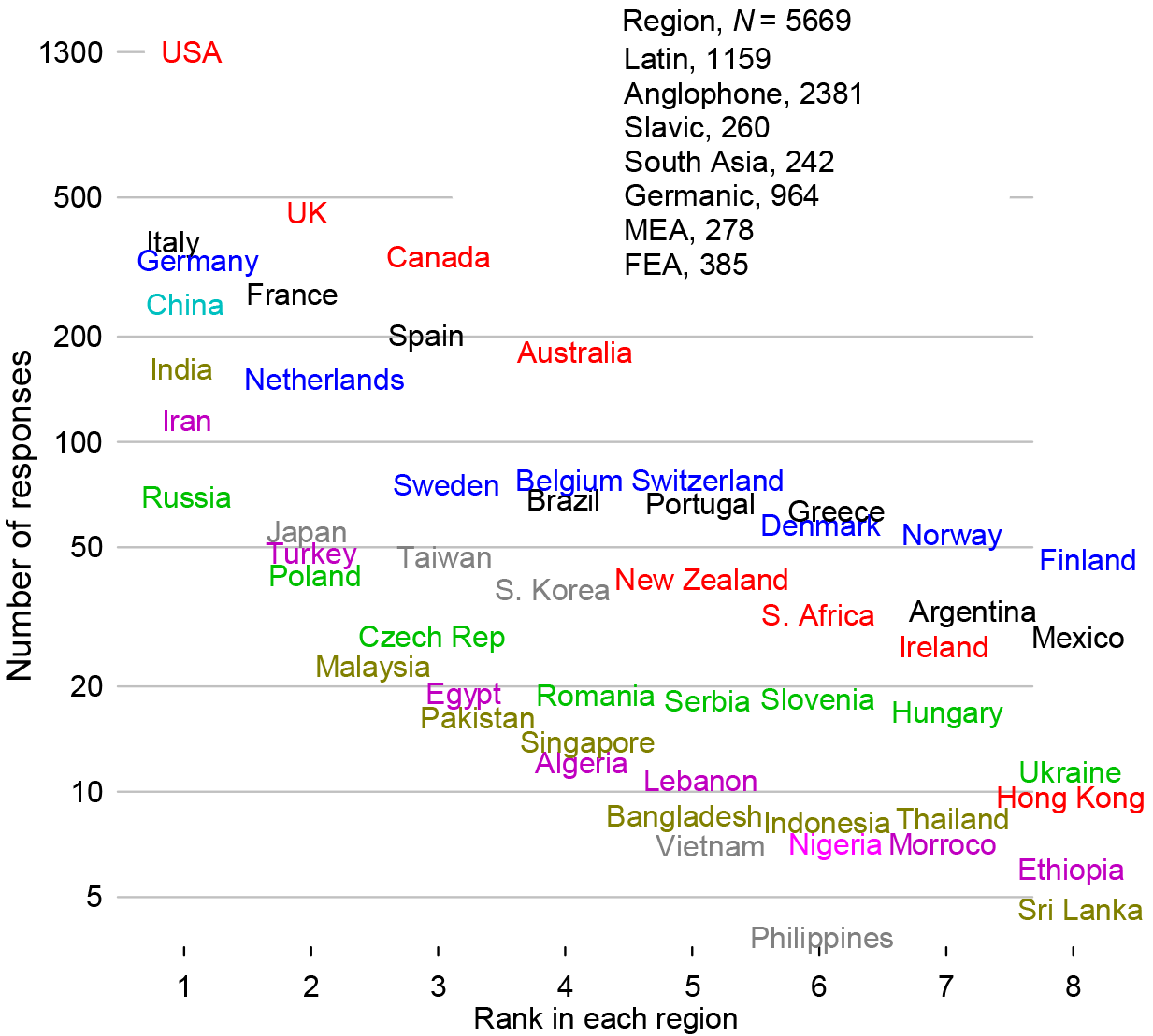
Number of respondents from each country and grouped in regions.

The differences between regions were much less than the differences between categories (Fig 8) and so birth place is an inconsequential contributor to the huge variance shown in Fig 3. Some slight differenes are evident: anglophones responded similarly to northern Europe (Germanic) each with *S̄*_*j*_ = 53, which means they are less generous assigning authorship than the other regions. Researchers from Far East Asia (FEA) and south Asia recognized non-standard contributions (that is, supervise, design, and execute) most (*S̄*_*j*_ = 62 and 61), while Anglophones and the germanic cluster recognized these least. Excluding sub-Saharan Africa and Iran, *S̄*_*j*_ = 62 for MEA. According to a *t*-test of the means, the responses from Israel were indistinguishable from the Germanic region. The responses from the Slavic group were much like the Latin group at *S̄*_*j*_ = 59. Correcting grammar is among the least valued activities but FEA and MEA researchers rank it close to a full point higher than researchers from other regions.

**Fig 8.**
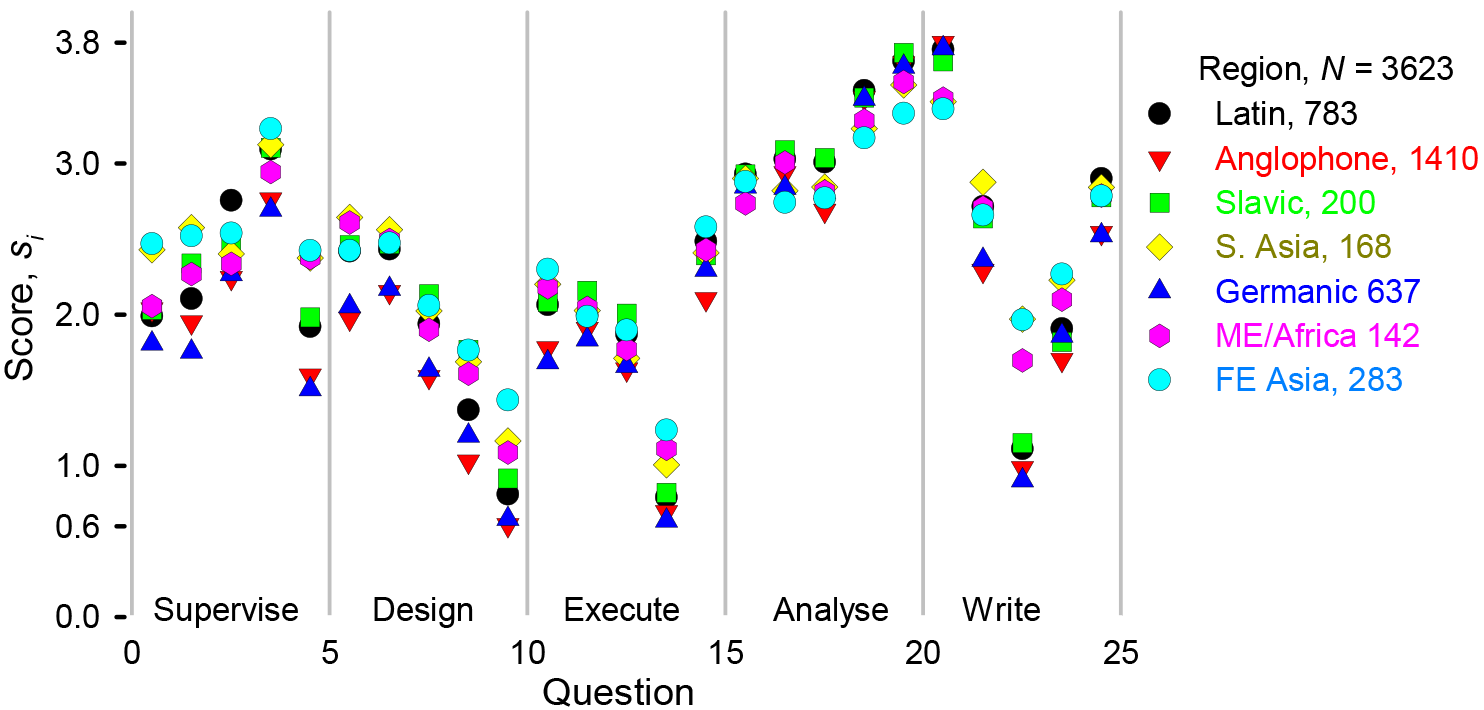
Activity scores as a function of region. Researchers from 115 countries answered the survey and we grouped them into regions based on *t*-tests comparing the mean score (*S̄*) of the country and region. The Latin region includes South American countries, Portugal, Spain, Italy, France, Greece, and Cyprus. Anglophones include the United States and many Commonwealth countries. Slavic nations comprise Eastern Europe and former Soviet states. India, Pakistan, Indonesia, Singapore, and Bangladesh are part of the South Asia region. The Germanic states include European countries north of France/Italy. Middle East and Africa (MEA) is the most diverse grouping and includes Iran, Turkey, Arabic-speaking nations and sub-Saharan Africa (some of which could have been included among the Commonwealth nations). China, Japan, South Korea, Taiwan, Vietnam comprise the Far East Asia region (FEA).

## Discussion

For almost every category, opinions range between the two extremes (Fig 3): as many people think that they should rarely ever merit authorship as those that think is should almost always merit it. Describing the data statistically and comparing responses across fields, countries, and profession identifies trends but the large variance show how divergent opinions are. Even within the same discipline, same region, and same level of experience, responses extended from one extreme to the other. These data demonstrate that the top cited authors disregard the ICJME critera, but how about the medical profession? The responses of the 722 researchers in medicine were indistinguishable from all other researchers Fig 9 with Fig 9: everyone continues to recognize contributions other than those outlined by the IJCME. The NIH rated how often activities in their guidelines merited authorship and we have reproduced their estimates in Fig 9 (NIH label). Their scores agree with the respondents except for commenting on the manuscript and providing resources with scores of 2.6 (*s̄*_22_) and 2.2 (*s̄*_1_), respectively.

**Fig 9.**
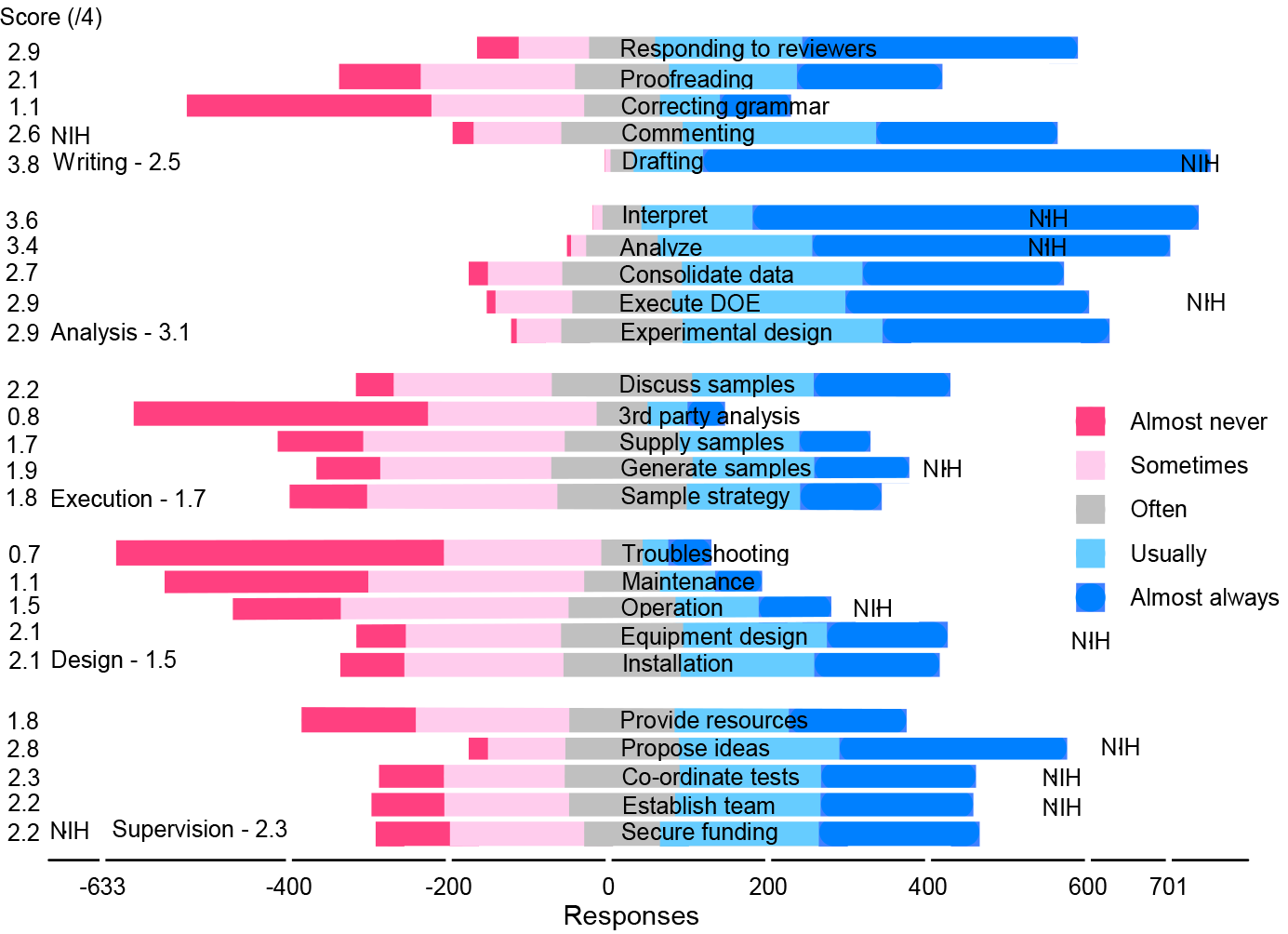
Medical sciences Likert chart: 720 respondents. The number of responses less than zero on the *x*-axis include almost never (*ω*_0_), sometimes (*ω*_1_) and half of the often (*ω*_2_) responses. The number of responses greater than zero include almost always (*ω*_4_), usually (*ω*_3_), and half of the often responses. The mean scores on the *y*-axis are *s̄*_*i*_ = Σ_*k*=1,*n_i_*_ ω_*i,k*_/*n*_*i*_ *s̄*_*i*_ and vary from a low of 0.7 (troubleshooting) to a high of 3.8 (drafting).

The ICMJE and NIH recommend how to attribute authorship but our survey demonstrates that the most successful researchers in the world recognize intellectual content beyond their criteria—research is messy and universal guidelines may not infringe on the autonomy of scientific practice [17, 28]. Assigning authorship relies on fairness on the part of principal investigators (PI) that receive public and private funding. They have a duty to conduct research responsibly, which comprises honesty, integrity, openness, and transparency. PIs have the authority to choose individuals and groups to conduct the work, the authors, and author order. However, PIs have an obligation to share their authorship criteria so that everyone understands how they will be recognized—authorship, acknowledgment, financial compensation, etc. A point system that assigns weights to research activities is one approach to assess contribution [29, 30]. The PI assesses input from each of the contributors and divides the points of each activity among them. Individuals can contribute to several activities and anyone that exceeds a predetermined threshold becomes an author.

Tangible research outputs include ideas, data, designs, writing, graphs, programs,and methodologies and these may be protected by copyright or patents. Researchers have the sole right to reproduce, distribute, publish, and create derivatives of their original work. As soon as work is recorded (written), publicly or privately, it is covered by copyright, which raises questions regarding how to acknowledge professional writers, particularly those with subject specific knowledge that correct and edit not only text but also improve scientific content [31]. Excluding creators of original work constitutes 338 copyright infringement, regardless of how few points they have accrued. Copyright does 339 not protect ideas but copying an idea, data (interpretation), methodologies could be plagiarism. The system of points is appropriate to assign author order but other considerations may trump it for attributing authorship.

## Conclusion

Researchers publish to build a reputation that universities and companies examine to hire and promote. Consequently, authorship lists are growing, so journals require everyone to disclose their contribution to ensure equitable recognition—authorship or acknowledgment. However, articles continue to include individuals with a modicum of intellectual involvement [32]. Our survey demonstrates that people value activities beyond writing and analysing data but the opinions are polarized: As many researchers credit activities like supervision almost always as those that almost never, even people in the same field and country. Pure and applied scientists grant authorship outside the ICMJE guidelines more compared to those in the humanities. This prevalence is related to large collaborations and the importance of building social relationships. The guidelines are a means for researchers to discuss social pressures regarding authorship but, adding individuals with imperceptible contributions is more tolerable than excluding those adding creative content.

